# Evaluating the Number of Different Genomes in a Metagenome by Means of the Compositional Spectra Approach

**DOI:** 10.1101/2020.07.23.217364

**Authors:** Valery Kirzhner, Dvora Toledano-Kitai, Zeev Volkovich

## Abstract

Determination of metagenome composition is still one of the most interesting problems of bioinformatics. It involves a wide range of mathematical methods, from probabilistic models of combinatorics to cluster analysis and pattern recognition techniques. The successful advance of rapid sequencing methods and fast and precise metagenome analysis will increase the diagnostic value of healthy or pathological human metagenomes. The article presents the theoretical foundations of the algorithm for calculating the number of different genomes in the medium under study. The approach is based on analysis of the compositional spectra of subsequently sequenced samples of the medium. Its essential feature is using random fluctuations in the bacteria number in different samples of the same metagenome. The possibility of effective implementation of the algorithm in the presence of data errors is also discussed. In the work, the algorithm of a metagenome evaluation is described, including the estimation of the genome number and the identification of the genomes with known compositional spectra. It should be emphasized that evaluating the genome number in a metagenome can be always helpful, regardless of the metagenome separation techniques, such as clustering the sequencing results or marker analysis.

## Introduction

In this paper, we present a method of counting the number of genomes in the test medium. The suggested approach is based on the compositional spectra (CS) method, proposed a long time ago [1-5] for the comparison of genomes and/or long genome fragments.

By definition [3], the *compositional spectrum* (CS) is the frequency distribution of oligonucleotides of length *k* (in the literature, referred to as words, *k*-grams, or *k*-mers) which occur in the genome sequence. The existing versions of the method differ mainly in the choice of the set of oligonucleotides, called *support* (*dictionary*), for which the frequency distribution is calculated. At present, there exists a large body of research on genome comparisons, which employ different versions of this method and produce results indicative of its validity (see, e.g., [6,7]).

Here the CS method is employed to analyze the result of metagenome sequencing, which represents a set of all words of fixed length composing the genomes of the metagenome, with regard to their multiplicity. Since the orientation of each word is unknown, we symmetrize this set by duplicating each word in two possible orientations. Then the set of the metagenome words can be represented as the union of CS^+^ and CS^−^ spectra of each genome (with regard to its multiplicity), which are calculated in both chains in the 5’-to-3’ direction. The union CS ^+^ ∪ CS ^−^ is referred to as the *barcode spectrum* (*BS*) [8]. For our purposes, *BS* is considered a vector, with each coordinate of the vector corresponding to a word from the chosen dictionary. Thus, the dimension of vector *N* grows with the growth of the number of words in the dictionary.

The calculations are based on the following statement [9]: “If the number of the genomes under consideration, *n*, is less than the space dimension, *N* (*n < N*), *there are no biologically significant reasons for the CS vector of one genome to be in the linear span of the CS vectors of the set of some other genomes”*. (If *n > N*, then, for purely formal reasons, *n*-*N* spectra are linear combinations of all the others.) The same is true for the barcode spectrum. For a more detailed description of these definitions, see [9,10].

There exist numerous *in silico* methods for separating the genomes constituting a metagenome that employ a k-mer set, obtained by the metagenome sequencing. In a large group of methods, all the segments constituting the mixture are partitioned into clusters in such a way that each cluster contains only the segments belonging to the same genome. Obviously, the result of this procedure is the *separation of the genome mixture*. These methods employ the distances between the segments, which are defined on the basis of different features of the segments such as C+G content, dinucleotide frequencies, and synonymous codons [11-13]. The segments may be also characterized by fixed-length words (see, e.g., [8], where the length of 4 is assumed). It should be noted that when the separation problem is formulated in such a way, the methods of its solution do not require—generally speaking—knowledge of the genome sequences of the bacteria constituting the mixture.

The methods of another group, which can be viewed as the opposite of the methods described above, check the presence of a particular microorganism genome in the mixture. Obviously, these methods can be applied only if the genome sequence (not necessarily the whole sequence) is known. The methods that are usually employed in this case search for the similarity between the known genome sequences and the fragments constituting the mixture. Some of these methods, based on the BLAST methodology, use marker genes [14], DNA-polymerase genes [14], and genes encoding protein families [15].

In the above-cited studies, the k-mers, as a rule, are united in a cluster corresponding to one genome on the basis of pairwise comparison. In contrast, in [16-18], the whole k-mer vector is considered as genome characteristics. In [16,18], the composition similarity for different metagenomes is assessed by comparing their k-mers vectors. In [17], the presence of a particular vector in the metagenome vector is assessed by machine learning methods.

The issue of the number of unknown genomes in a metagenome, investigated in our work, had already been considered (see, for example, [19, 20]). These studies employed procedures which could be used only for one sample of the metagenome. In contrast to this, our approach is based on analyzing the compositional spectra of subsequent samples of the medium. Its essential feature is using random fluctuations in the number of bacteria in different samples a metagenome. The idea of using several samples from the same metagenome was suggested in [21], where its effectiveness was proved for analyzing certain metagenome parameters.

In the “Calculation of the number of different genomes in the medium” Section, the concept of a *sample* is introduced as a metagenome extracted from the medium under study. It is essential that, due to the random character of the samples, they—generally speaking—contain a different number of genome copies (have different multiplicity). It is shown that the number of different genomes in the test medium can be calculated by sequentially sampling the medium. The content of this section is covered in [22].

The possibility to effectively distinguish between these two states of a sample set is analyzed in the “Computational aspects of the proposed approach” Section.

## Results

### Calculation of the number of different genomes in the medium Set of the samples of the medium

Let us refer to a certain portion of the metagenome as the *parent population*. We define the *metagenome samples* as random samples from the parent population. For further consideration, it is essential that the number of any genome copies in different samples is a random value, independent of those for other genomes. This follows from the statistical laws of the “parent population – sample” relation. The distribution for these random values depends on the properties of the test medium and on the relative volume of the samples. In the present work, we assume that the samples are of equal volume and that the general population volume is large enough as compared to the sample volume for the multiplicities of the genomes constituting the general population, being practically independent of sampling without replacement. Then the mean value of any genome multiplicity over all samples is proportional to this genome multiplicity in the general population, the coefficient of proportionality being the same for all the genomes.

Consider the properties of sequence *P* = {*p*_1_, *p*_2_, …} of independent samples of the medium (where ***p***_*i*_ is the *i*^th^ sample). Let *T*_*i*_ be the *vector of genome multiplicities (VGM)* in the *i*^th^ sample, ***T***_***i***_ = (***t***_***i***1_, ***t***_***i*** 2_, …, ***t***_***in***_), where *t*_*ij*_ is the number of copies of the *j*^th^ genome in the *i*^th^ sample. Below we present the two most probable models of genome occurrences in the sample in terms of VGM.

<u>***Model 1***</u>. It is assumed that the number of each genome copies in a sample has normal distributionThen, the number of copies of the *j*^th^ genome in each sample can take any value out of set {*m*_*j* ±_*μ*_*j*_} (*μ*_*j*_ = 0, 1, 2,…, *s*-1), where it is naturally supposed that *μ*_*j*_ <*m*_*j*_.. The value of *m*_*j*_ depends on the number of genome *j* copies in the general population. Suppose further that each genome multiplicity in the sample is independent of the multiplicities of other genomes in the same sample. Then VGM vectors take the values out of set

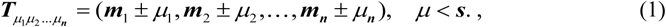

where multi-index *μ*_1_*μ*_2_ …*μ*_***n***_ determines each vector of set (1). The values of *μ*_*_ are independent of each other and take the values out of set {0, 1, 2, …, *s*-1}.

<u>***Model 2.***</u> It is assumed that the number of each genome copies in a sample has uniform distribution. It can be supposed that the *i*^th^ coordinate of a multiplicity vector *T*, being independent of the other coordinates, can take values out of a finite set of non-negative integers, *a*_*i*_={*a*_*ij*_}, *j*=1, …, *s*. Then the VGM set consists of all vectors of the type

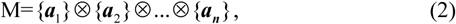

where ⊗ designates a direct product and

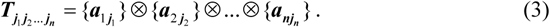

### Basic procedure

The barcode spectrum of each sample is a vector in N-dimensional vector space, which is a linear combination of BS spectra of some genomes of the medium. Thus, the number of samples that have linearly independent spectra is equal to the number of different genomes (i.e., without regard for their multiplicities) in the medium, which is denoted above by *n*.

Now let us sequentially take samples out of the medium and use them to build the basis of the sample BS spectra. This basis (the basic set) certainly contains the spectrum of the first sample. The spectrum of each consequent sample belongs to the basic set if it is linearly independent of the spectra of already chosen samples. In this way, by sequentially taking the medium samples, it is possible to obtain the full basis for the medium so that none of the further taken samples gives a new independent spectrum. The size of the obtained basis is equal to the number of genomes with linearly independent spectra that are present in the medium.

### Statistical estimations for the procedure of building the full sample basis Estimation of the fractions of the sample set inside and outside the plane

If vectors are sequentially and randomly taken out of an *n*-dimensional space with a standard measure, the first chosen *n* vectors constitute the basis of the whole space with probability 1. Indeed, if the first *p* vectors chosen in this way are linearly independent, then their linear span is a set of zero measure in the whole space. Therefore, the probability of choosing the next vector from this set is zero. However, if the measure is concentrated on a finite set, the above description is incorrect. Indeed, it is sufficient to notice that in this case a point can be randomly chosen twice with non-zero probability

Consider the general scheme described in Model 2.

#### Proposition 1.

In *n*–dimensional space, the fraction of the points of set *M* (defined by (2)) that lie outside any plane *L*_*p*_ of codimension *p* (0 < *p* < *n*) is not less than 1 − ***s***^− ***p***^.

**Proof**. Let us prove **Proposition** by the method of induction on the space dimension, *t*. Namely, the plane of codimension *p* appears when the space dimension *t*=*p*; i.e., in *p-* dimensional space, and has zero dimension, thus being a point in space *t*=*p*. In this space, set *M* consists of s^t^ points *M*_*(t=p)*_={***a***_1 ***j***_}⊗{***a***_2 ***j***_}⊗… ⊗{***a***_***tj***_}, with not more than one of the points lying on the plane of zero dimension. Consequently, the fraction of the points that lies outside this plane is (s^t^ -1)/s^t^, which equals 1− ***s***^−***p***^ at *t*=*p*, as stated in **Proposition**. Now, let **Proposition** be true for some dimension *t*. Let us show that it is also true for dimension *t*+1. In the space of dimension *t*+1, set *M* consists of points *M*_*t+*1_= {***a***_1 ***j***_}⊗{***a***_2 ***j***_}⊗… ⊗{***a***_***t*** +1, ***j***_}. We can present this set in the form of a union of the planes:

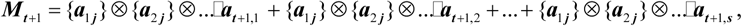

expanding *M*_*t+*1_ over the last coordinate. Planes

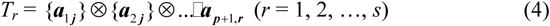

in this representation are, obviously, *t-*dimensional; the dimension of plane *L*_*p*_ in the space of dimension *t + 1* being equal to *t+1*-*p*. There exist two main cases of the dispositions of planes *T*_*r*_ and *L*_*p*_ with respect to one another:

(1) At *p* =1, one of the planes *T*_*r*_ may coincide with plane *L*_*p*_, whose dimension, in this case, is also *t*. However, the vertexes of all the other planes (4) will lie outside plane *L*_*p*_. Since, in *t+1*-dimensional space, the total number of *T*_*r*_ planes is s^t+1^, and in each plane there lie *s*^*t*^ vertexes, the fraction of the vertexes lying outside *L*_*p*_ is (*s*^*t*+1^ *s*^*t*^)/*s*^*t*+1^; i.e., (1-*s*^-1^). The latter expression coincides with the one in **Proposition** at *p*=1. It should be noted that in this case it is not necessary to make the suggestion of induction.

(2) Let plane *L*_*p*_ of dimension *t+1*-*p* intersect with plane *T*_*r*_ of dimension *t* for some value of *r* (*r* = 1, 2, …, *s*). Then the plane of intersection, *L*_*p,r*_, has a dimension of *t*-*p*; i.e., its codimension in plane *T*_*r*_ is *p*. Under the suggestion of induction, it follows that the fraction of points M in plane *T*_*r*_, which lie outside plane *L*_*p,r*_, is more than (1-s^-p^). Without loss of generality, it can be suggested that plane *L*_*p*_ intersects with all planes (4). (Obviously, this suggestion can lead only to the reduction of the number of points lying outside plane *L*_*p*_.) Since planes (4) have no common points and the fraction of the points lying outside plane *L*_*p*_ is not less than (1-*s*^*-p*^) for all planes *T*_*r*_, the same result of fraction (1-*s*^*-p*^) will be obtained *M*_*t*+1_, which proves **Proposition**.

If the random choice of a sample has uniform probability, then **Proposition 1** has an important

#### Corollary.

If, among the already-chosen samples, there exist *p* linearly independent ones, the probability of the next random sample being linearly independent (if such a sample still exists) equals 1− ***s***− ***p***. The probability reaches its minimum, 1− 0.5− ***p***, at *s*=2. In particular, in the case of the plane of codimension 1 (hyperplane), not less than 0.5 points of set *M* (samples) remain outside this hyperplane.

According to **Proposition 1**, the probability of finding the next linearly independent sample is a function of the dimension of the already found basis vectors with respect to the full dimension of the sample set. Since the latter parameter is unknown (the evaluation of the full dimension being just the goal of the study), at each step we should choose the probability corresponding to codimension 1. Obviously, this probability value is the minimal of all codimensions, thus the estimation is correct.

Now consider the situation where some probability measure is preset over the set of all possible samples. In the general case (within the framework of Model 2), let us arrange the number of genomes corresponding to each coordinate in the order of descending probability of finding a sample with the number of genomes ***p***(***a***_***i***1_) ≥ ***p***(***a***_***i*** 2_) ≥… ***p***(***a***_***is***_) (*i*=1, 2, …, *n*). Next, define the probability of each VGM, *P*(*T*), as the product of the probabilities of its coordinates. According to Proposition 1, the number of VGMs lying outside a plane of any codimension is not less than the number of VGMs lying in this plane. However, their total probability measure may now be less than 0.5 and a different estimation is required. For an algorithmic estimation, there is no need to evaluate the total probability value for all codimensions. As mentioned above, it is enough to obtain the estimation for codimension 1.

#### Proposition 2.

If the probability distribution results in inequalities ***p***(***a***_***i***1_) ≥ ***p***(***a***_***i*** 2_) ≥… ***p***(***a***_***is***_), the probability measure of all the vectors of set *M* lying outside hyperplane *L*_1_ of codimension 1 is not less than

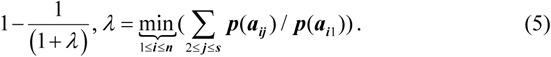

**Proof**. Outside hyperplane *L*_1_, there necessarily lies at least one unit vector *e*_*i*_, whose coordinates are all zero and only coordinate *i* equals 1. Otherwise, if all unit vectors lie in the hyperplane, its dimension coincides with the dimension of the whole space. Consider vector *T* belonging to the set of genome multiplicities, *M*, and lying in hyperplane *L*_1_. The value of the *i*^th^ coordinate of vector *T, a*_*iq*_ has probability *p*(*a*_*iq*_), where *q* is some integer between 1 and *s*. Then, obviously, vector ***T*** − ***p***(***a***_***iu***_) / ***p***(***a***_***iq***_)***e***_***i***_ (***u*** ≠ ***q***) also belongs to set *M*, but lies outside plane *L*_1_. By definition, its probability is equal to ***p***(***a***_***iu***_) / ***p***(***a***_***iq***_)***P***(***T***). Thus, each vector *T* of set *M* that lies in plane *L*_1_ can be associated with *s*-1 vectors that lie out of this plane. This association is unique in the sense that, for each pair of vectors ***T***_1_ ≠ ***T***_2_ (***T***_1_, ***T***_2_ ∈ ***L***_1_), equation ***T***_1_ −*α*_1_***e***_***i***_ = ***T***_2_ −*α*_2_***e***_***i***_ is not possible. Furthermore, if *q*>1, also for *u*=1 ratio ***p***(***a***_***iu***_) / ***p***(***a***_***iq***_) is more than unit on the strength of the suggestion about the ordered probabilities (see above). As a result, the measure of some vectors ***T*** − ***p***(***a***_***iu***_) / ***p***(***a***_***iq***_)***e***_***i***_ (***u*** ≠ ***q***) lying outside plane *L*_1_ will be larger than the measure of the original vector *T* in plane *L*_1_ (***p***(***a***_***iu***_) / ***p***(***a***_***iq***_)***P***(***T***) > ***P***(***T***). To estimate the minimum of the ratio ***p***(***a***_***iu***_) / ***p***(***a***_***iq***_), we assume that at coordinate *i*, index *q*=1 for all vectors *T* lying in plane *L*_1_. Then, for index u=2, ratio ***p***(***a***_***i*** 2_) / ***p***(***a***_***i***1_) will be the largest of all possible ones, descending order ***p***(***a***_***i*** 3_) / ***p***(***a***_***i***1_) ≥ ***p***(***a***_***i*** 4_) / ***p***(***a***_***i***1_) ≥… holding to the last value of *s*. Thus, if the total measure of vectors belonging to set *M* and lying in plane *L*_1_ is equal to ***P***(***L***_1_), it will obey inequality 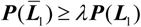, where 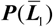 is the full measure for the vectors lying out of the plane, 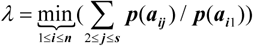. The ratio of the measure of the vectors lying outside hyperplane *L*_1_ to the measure of all the vectors of set *M* is 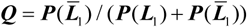, which results in ***Q*** > *λ****P***(***L***) / (***P***(***L***) + *λ****P***(***L***)). Thus, ***Q*** > *λ*/ (1+ *λ*) = 1−1 / (1+ *λ*). This estimation coincides with the one obtained in **Proposition 1** for the uniform measure multiplicity, ***p***(***a***_***i***1_) = ***p***(***a***_***i*** 2_) =… = ***p***(***a***_***is***_), because in this case *λ*=*s*-1.

In Model 1, in the case of normal distribution of genome multiplicities, the probability of the genome multiplicity value at coordinate *i* is

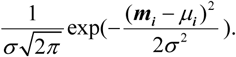

According to (5), *λ*= exp(−1 / 2σ_0_^2^), where σ_0_ =min(σ_i_), *i*=1, 2, …, *n*. For large σ_0_, the value of *λ* approaches 1 (e.g., at σ_0_=10, λ=0.995).

Next, we evaluate the mean number of samples that must be taken to obtain all the basis vectors of the sample set.

#### Proposition 3.

The mean number of samples required to find all *n* basis vectors is equal to *n*/*p*, where *p* is the probability of finding a basis sample in one operation.

**Proof**. The probability of finding the *n*^th^ basis element in the (*n + t*)^th^ sampling is

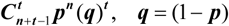

(according to [23] I, VI, 2). Then the mean length of the sampling sequence up to the step of finding all *n* basis elements is

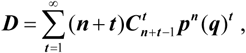

or

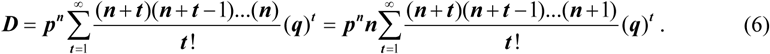

It is easy to show that

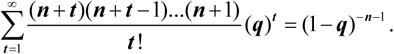

Substituting the latter expression into (6), we obtain ***D*** = ***pn*^*n*^**(1− ***q***)^−***n***−1^ = ***np***^−1^, which proves **Proposition 3**.

In the case of uniform distribution considered in **Proposition 1**, *p* = 0.5, and the mean value of samples is 2*n;* while, according to **Proposition 2**, for any measure used in it, the mean value of samples is (1+ *λ*)***n*** / *λ*.

The last step in building the above algorithm is formulation of the rule of its termination. It follows from the above that, if in *m* consequent samplings, the spectra are found to be linearly dependent on the already chosen basic set, then e.g., in the case of uniform distribution of the samples, the probability of all the basis samples having been already found is 2^-*m*^. For example, at *m*=4, it can be claimed with an accuracy of 94% that there are no more independent samples.

### Computer simulation of the fractions of a sample set inside and outside a plane

In the previous section, the parts of the set of samples inside and outside the plane (hyperplane) are estimated. These estimates are accurate in the sense that it is possible to give examples when the estimates cannot be improved. For example, consider the set of all metagenome samples, *M*, in Model 2, provided that in each sample the number of copies of the *j*-th genome can take values from set {0 _±_*μ*_*j*_} (*μ*_*j*_= 0,1). If random sampling of *n-*1 samples is performed in *n*-dimensional space, it may happen that these samples are unit vectors ***e***_1_, ***e***_2_, …, ***e***_***n***−1_. Then in the corresponding hyperplane, there exist, in total, 2^n-1^ elements of set *M*, and outside this hyperplane there also exist, in total, 2^n-1^ elements. Thus, the probability of randomly choosing the next sample from set *M* lying partly outside the hyperplane is 0.5, which is in line with **Proposition 1** (see Corollary to **Proposition 1**).

On the other hand, if randomly selected *n-*1 samples have the form 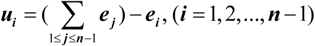, then in such a hyperplane there exists no vector from set *M* that is different from basis *u*. In this case, inside and outside the hyperplane, there lie *n-*1 and the remaining 2^n^ – (*n-*1) vectors from set *M*, respectively. Thus, the probability of randomly choosing the next sample from set *M* part lying outside the hyperplane is much larger than 0.5 and rapidly converges to 1 with increasing *n*. This example shows that the real probability of choosing a linearly independent sample (under the assumption that such a sample exists) can be much larger than the theoretical estimates.

This issue was investigated by computer simulation. Fig.1 shows the probability of *n* samples randomly chosen from a metagenome composed of *n* different genomes having linearly dependent BS vectors. To calculate each point of the curves, one million trials were done. The obtained curves fit the following expression perfectly:

**Figure 1.**
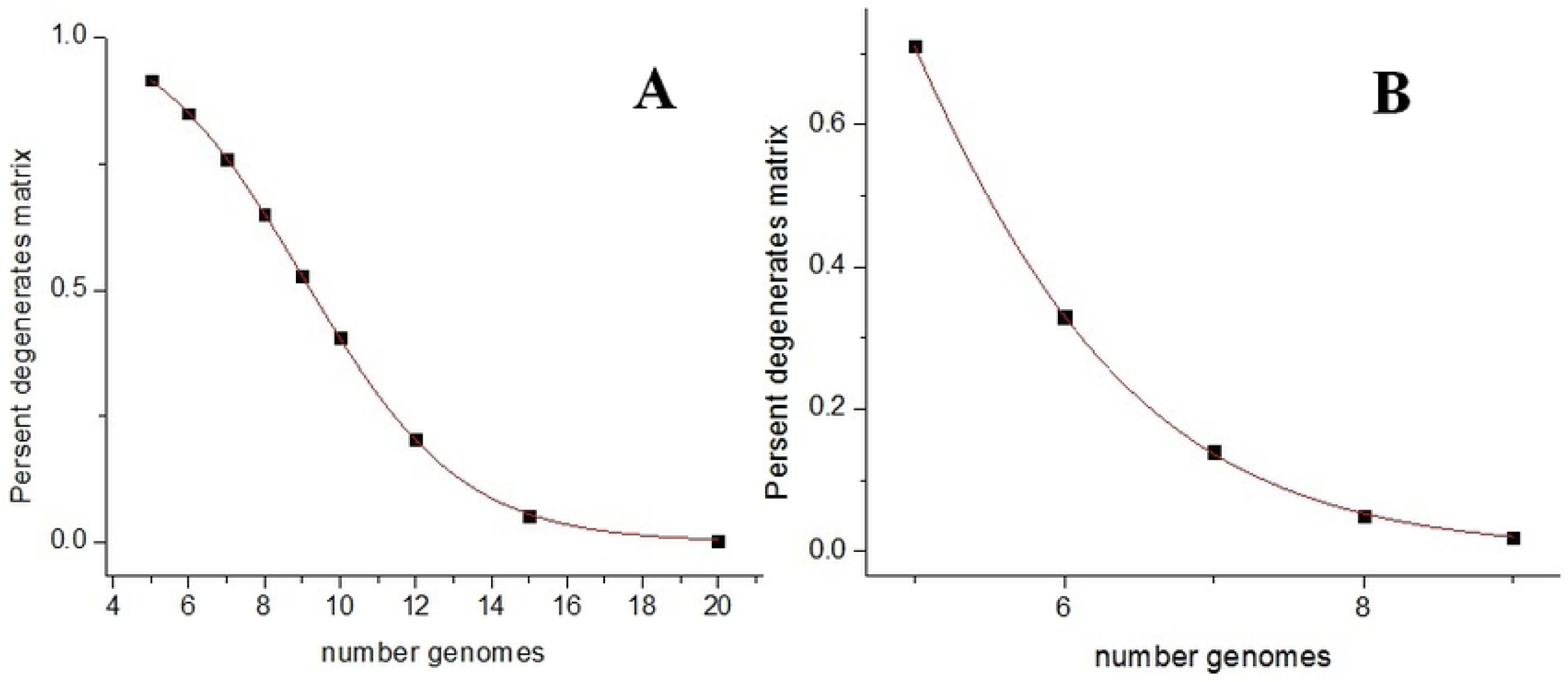
Probability of *n* samples randomly chosen from a metagenome composed of *n* different genomes being linearly dependent. (A) Each genome is present in the sample with equiprobable multiplicity of 0 or 1. (B) The same for four equiprobable multiplicities (0, 1, 2, and 3). *x*-axis: the number of different genomes in the metagenome; *y*-axis: percentage of linearly dependent samples.

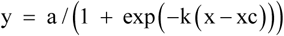

For the two curves, parameters *a, k*, and *c* are different. It can be seen that the increase of the volume of a sample set results in the almost unit probability of the randomly chosen samples being linearly independent if their number does not exceed the basis of the sample space. Thus, the discrete set of sample vectors begins to behave like a continuous set with the increase of the number of genomes in the metagenome.

### Computational aspects of the proposed approach

In this Section, we consider the fundamental possibility of calculating the number of different genomes in the metagenome in the presence of data errors.

#### Design of the numerical experiments

In each experiment, the number of genomes in a metagenome is numerically determined based on a set of simulated samples of this metagenome. For the metagenome, number *p*, the number of different genomes in the metagenome, is specified, and list *P* of these genomes is randomly selected from library *L* [9], containing 100 bacterial genomes. Then *N* samples are formed, each including all genomes of list P with multiplicities that have normal distribution (m, σ) identical for all genomes. The BS spectrum of a sample is the sum of the BS spectra of all its genomes, possible errors in their sequencing being taken into account. Namely, each genome g from library L is assigned set G (g, 75), which contains twenty randomly selected fragments, each comprising 75% of total genome g (25% error). Each fragment is also transformed by randomly replacing 1 out of 1000 letters [24]. The BS spectra of these fragments are used for the formation of the sample spectrum as follows: if a given sample contains *m* _*_ copies of genome g, then *m* _*_ more fragment spectra from set G(g,75), chosen with equal probability independently of each other, are added to the sample spectrum. In this way, errors in the determination of letters and the loss of parts of the genome during sequencing are simulated. In the study full 6-mers spectra are used.

In the course of experiment, number N of samples taken from the metagenome increases by one to a predetermined boundary. For each N value, the number of different genomes is calculated. Such experiment is repeated 100 times, each time with a different random list of P genomes. As a result, we obtain the dependence of the percentage of correct answers on the number of samples used.

We also use set G (g, 80) of twenty fragments, each containing 80% of the full genome and set G (g, 90) of 10 fragments, each containing 90% of the total genome. In the latter case, each genomic sequence is divided into 10 equal non-overlapping parts and 10 spectra are calculated, each spectrum excluding one of the parts. Thus the obtained spectra correspond to 10% of data errors, while their pair-wise difference is 20%. In all these cases, single letters are also changed according to the described above scheme.

At the end of this section, we estimate the magnitude of the error introduced into the 6-mers spectra by replacing complete genomes by their parts. The corresponding vectors (BS spectra) should not be collinear, but how large are the angles between them? Fig. 2 shows the distribution of pairwise angles between 6-mers spectra of genomes from library L (Fig. 2 D) and of pairwise angles between the spectra of genome fragments comprising 75% and 50% of the complete genomes (Fig. 2 C and B, respectively).

**Figure 2.**
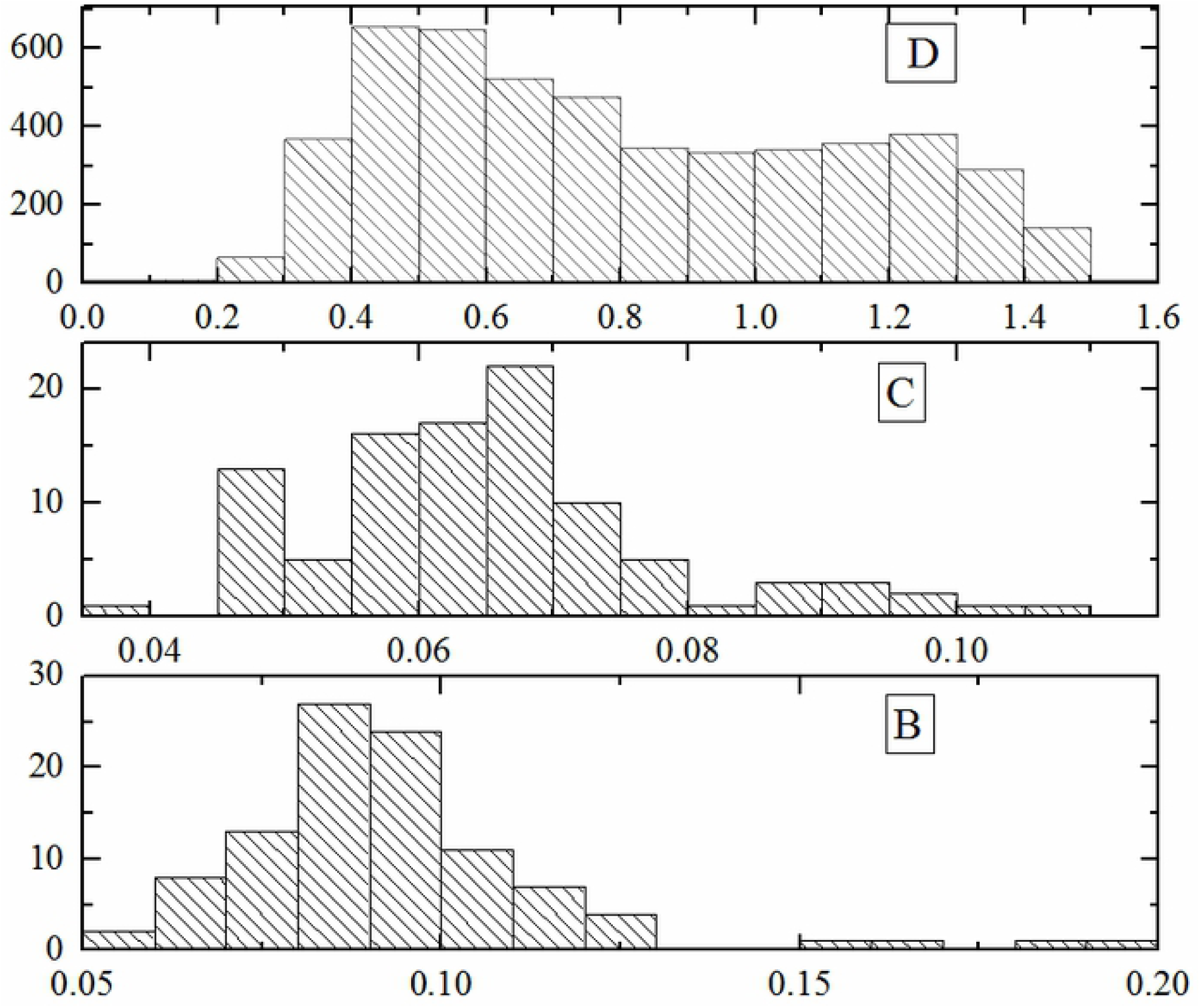
Distribution of pairwise angles between the BS spectra of complete genomes and between the BS spectra of the same genome fragments. (D) All pairs of complete genomes from library L. (C), (B) All pairs of the same genome fragments for all genomes from library L; the fragment sizes are equal to 75% (C) and to 50% (B) of the corresponding total genome. X axis: angles in radians.

It can be seen that even in case B the angles are much smaller than in the case D of complete genomes. It can be seen that data errors, even as high as in case B, still do not prevent the evaluation of the number of different genomes. It was shown by us earlier [3,25], that a fragment spectrum usually almost coincides with the whole-genome spectrum for a large number of genomes. The exception from this finding is a small number of certain bacteria that have significant differences in the spectra of their genome fragments, namely, *Borrelia burgdorferi* [3, 26], and some *Spirochete* bacteria.

### Computational method

In the absence of data errors, the dimension of any set of BS sample spectra does not exceed the number of different genomes in the metagenome under study, and all samples lie in linear subspace T, whose dimension is equal to the number of different genomes in the metagenome. By sequentially increasing the number of samples and calculating, at each step, the rank of the set of their spectra, we can calculate the dimension of subspace T, and thus obtain the number of different genomes in the metagenome, as described in the theoretical part of the work. In the presence of data errors, the dimension of the set of sample spectra is, generally speaking, equal to the number of samples. However, all the BS vectors lie near to plane T and the solution is equal to the dimension of this plane. The proper technique for solving such a problem is the principal component analysis (PCA) [27]. According to this method, sequential planes (linear subspaces) are constructed on the vectors of the main components, obtained on the basis of a matrix whose columns are the sample spectra vectors and then the total distance of the latter vectors from the corresponding planes is calculated. This result will decrease monotonously with each subsequent plane, but it can reach zero only in the absence of data errors. Therefore, it is necessary to find a certain “marker pattern” which would appear in the course the described process, signalizing that the dimension of the next subspace is equal to the number of different genomes in the metagenome.

We found such marker pattern, studying the characteristics of the method of principal components for various values of the sample parameters. Consider a multidimensional plane built on the first n principal vectors numbered in their standard order. Denote by f (n) the sum of the distances of the vectors of all the samples under consideration from this plane. Fig. 3A shows the dependence of f(n) on n. This curve, indeed, decreases monotonously. Fig. 3B shows the “normalized derivative” of f(n):

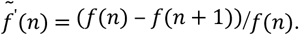

**Figure 3.**
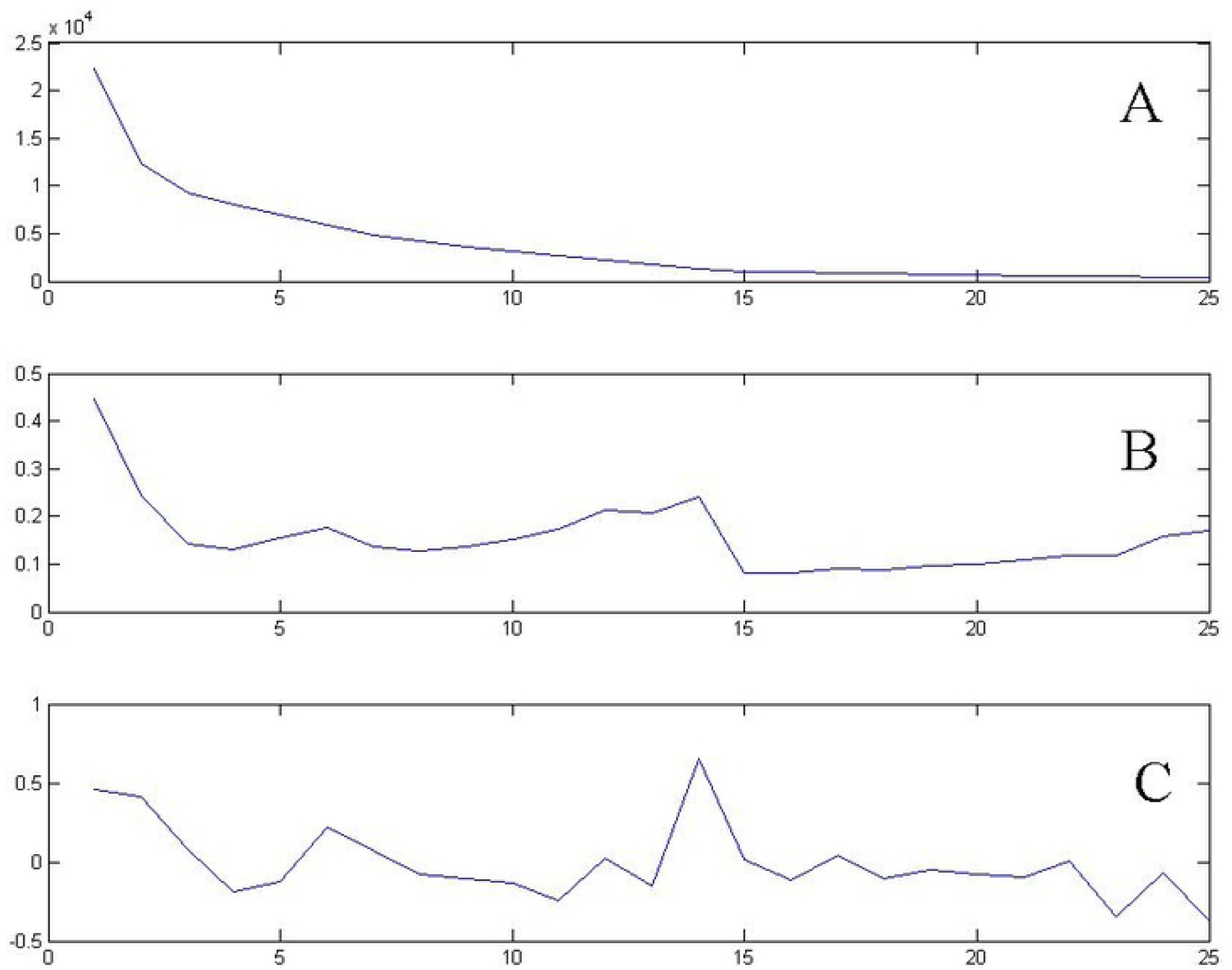
Dependences of the total distance between all considered sample vectors and multidimensional planes built on the sets of the corresponding principal components (A). The same for the normalized values of the first (B) and the second (C) derivatives of function (A). The metagenome contained 15 different genomes and 30 samples were used at m = 10, σ = 3. X axis: the number of the first principal components on which the plane is built.

One can see a specific step of curve 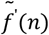 at n = 14-15. Fig. 3C shows the “normalized second derivative” of f (n):

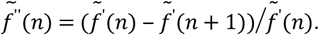

It can be clearly seen that curve 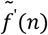 in Fig. 3B sharply decreases at n = 14 or 15. This means that further addition of the main components has little effect on the change of f(n). Obviously, such pattern of curve 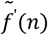 reflects the fact that the construction of the plane that best approximates the sample vectors has already been achieved.

The step-shaped pattern of the curve in Fig. 3B or the peak-shaped pattern of the curve in Fig. 3C were used to calculate the number of different genomes in a metagenome.

It should be also noted that the step-shaped pattern in Fig. 3B equally corresponds to the dimensions of 14 or 15, and the peak-shaped pattern in Fig. 3C corresponds rather to the dimension of 14, which is the correct value because while calculating the principal components, the set of sample vectors was shifted by the average value of the sum of all sample vectors, i.e., lost one dimension.

### Results of numerical experiments

In this section, we present some results of numerous experiments performed to estimate of the quality of the method. MATLAB was employed in this work, in particular, the main components were calculated using the princomp function.

The number of samples was consistently increased (as it would be done in a real-life experiment) and the quality of the simulation result was estimated. Computational difficulties in a real-life experiment do not exist, while in the case of numerous computational experiments the difficulty is coursed by a complicated nature of the marker pattern. Its correct automatic recognition being a separate problem, in this work we used a rather primitive method of classification, based on the “second derivative” maximum (see Fig. 3C). Although the results can be significantly improved with more accurate (even visual) pattern recognition, the computer simulation performed makes it possible to assess the proposed method for calculating the dimension of sample vectors using the principal component method and the method for estimating this dimension by means of the marker pattern.

#### Recognition of the linear dimension of a sample set for sequential increase of the number of samples

In these experiments, we used the value of data error of 25% in each genome, two values of the number of genomes in the metagenome (10 and 15), two values of the average concentrations of each genome (m = 10 and 20), and three normal distributions (m, σ) for σ = 1, 2, 3. It should be noted that the calculated mathematical expectation of random concentration deviation from average value *m* for the values of σ = 1, 2, 3 was 0.77, 1.58, 2.38, respectively. During each experiment, 50 samples were generated with the same composition of genomes, but different genome multiplicities (different number of genome copies) in each sample. For each genome g, the multiplicity value was determined by the normal distribution (m, σ) and then each copy of genome g was represented by the BS spectrum of a randomly selected fragment from set G (g, 75). Starting with five samples, we added one more sample from their remaining number in random order and, at each step, the described-above computational procedure was carried out to detect the appearance of a marker pattern. Such experiment was repeated 100 times. The accurate number of different genomes being known, we can compare the result of each single experiment with the number of samples used and thus obtain the reliability of the method.

The results of the computer simulations are shown in Fig 4. It can be seen that the reliability decreases with the increase of the number of genomes in a metagenome and the average number of genome copies. On the other hand, the greater is the scatter of the genome copies in the samples relative to the average, the better is the result. For example, if the metagenome contains 10 different genomes and the average number, m, of each genome copies is 10 at σ = 3, then only 20 samples are needed to evaluate the correct number of different genomes almost accurately (the error may be not more than ± 1). If the number of different genomes is 15, the number of copies being 10, the result is the same - the number of samples needed to obtain the correct number of different genomes almost accurately is 30, being also equal to twice the number of different genomes.

**Figure 4.**
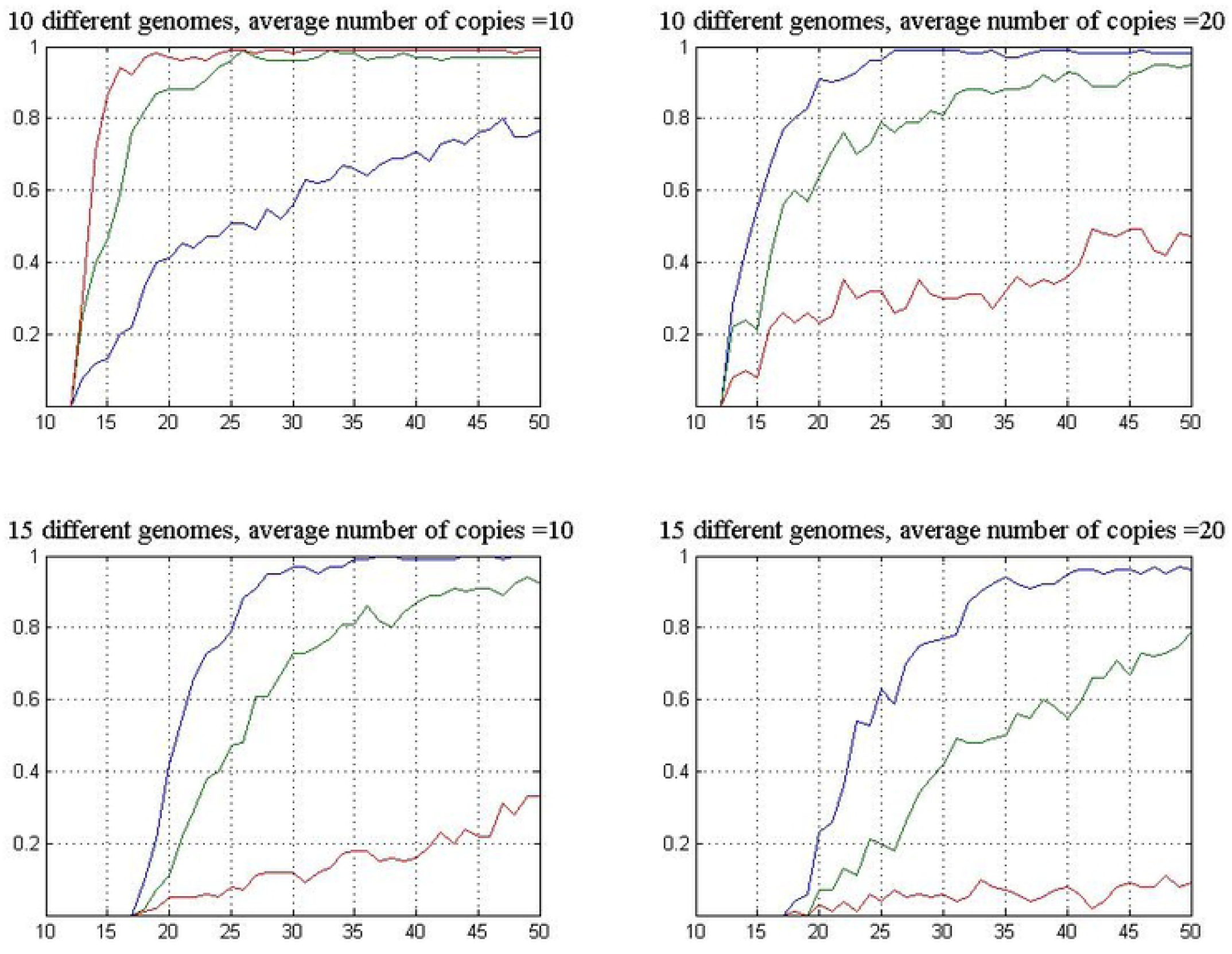
Results of estimating the number of genomes in a metagenome in the course of sequential increase of the number of samples. The BS spectra of genomes were taken from the corresponding fragment sets G (g, 75). In each chart, the lower, middle, and upper curves correspond to σ = 1, 2, 3, respectively. X axis: the number of samples; Y axis: the proportion of correct (the error not more than ± 1) to all results.

The curves in Fig. 5 show that the probability of the correct result increases with the decrease of the data error, at the same values of the metagenome parameters. At the error of 10%, only the curves for the worst results shown in Fig. 4 (10 and 15 different genomes at m = 20 and σ = 1) are presented in Fig. 5. Obviously, the reliability of the method is quite high and, obviously, it will be still higher with the increase of σ.

**Figure 5.**
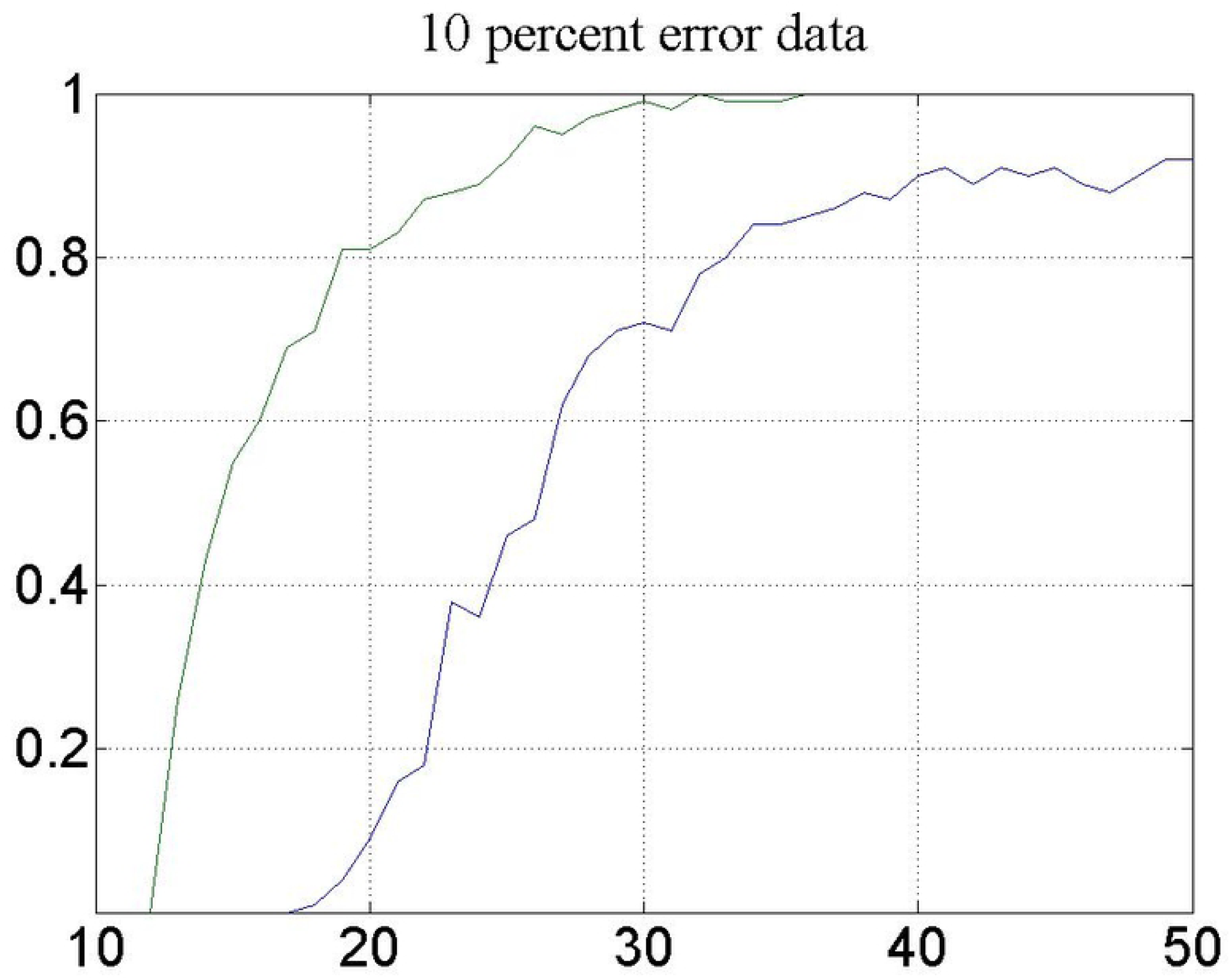
The same as in Fig 4, for fragment sets G (g, 90), i.e., at the data error of 10%. The average number of genome copies in the metagenome is 20; σ = 1. The lower and the upper curves show the percentage of correct answers at the number of different genomes 15 and 10, respectively.

The curves in Fig. 4 show that, for any metagenome parameters, there is a critical number of samples which provide high probability of the correct result obtained on the basis of the marker pattern. This critical value can be easily estimated in the course of simulation experiments. Since it is just the parameters of the metagenome in a real-life experiment that are unknown, the practical significance of the above result consists in the fact that the marker pattern indicates the right value of the linear dimension of a sample set even after passing the critical value of the number of samples used (Fig. 4). Such a “stationary” response to sequential addition of samples means the achievement of the critical value.

#### Recognition of the linear dimension of a sample set using already known genomes

If no information about the genomes that make up the metagenome is used, their number is estimated by the algorithm described above. However, the same method can be applied to a formally more general question, namely, about the number of different genomes in a particular metagenome that are not included in the given library of genomes represented by their BS spectra. If the library is empty, the problem appears to be the same as the one formulated above. If the library is not empty, its BS spectra should be used in the framework of the proposed method. The most obvious (but not the only one) way to do this is to form “virtual sample” spectra based on the library BS spectra. Then the real samples (i.e., those obtained from the metagenome under study) are combined with the virtual samples and the previously proposed algorithm is applied to the total sample set in the following way.

Suppose there are p different genomes in the metagenome. Part of these genomes (p_1_) are taken from the library and are combined with p_2_ genomes from the same library which are not present in the metagenome under study (but this is not known to us *a priori*). Virtual samples are formed from the BS spectra of these selected genomes in the ways described below. Applying our algorithm to the whole set of real and virtual samples, we will obtain that the total number of different genomes in the whole sample set is Z = p + p_2_. Since the virtual samples contain p_1_ + p_2_ different genomes, the number of unknown genomes can calculated as Z − (p_1_ + p_2_) = p − p_1_. This value is the number of different genomes in the given metagenome minus the number of genomes that were taken from the library and included in this metagenome. Thus the number of different genomes in the metagenome which are not included in the library can be obtained, but it is not clear which genomes from the library are included in this metagenome. Below a few possible ways of virtual sample design are considered.

1. Each virtual sample contains the BS spectrum of one genome from the library.
2. Each virtual sample contains a random mixture of BS spectra of the genomes selected from the library.
3. Each virtual sample contains the BS spectrum of one genome from the library, in which a random error is introduced.
4. Each virtual sample contains a random mixture of BS spectra of the genomes selected from the library, each genome containing a random error.

If only linear dependencies were considered, then the above first two ways of virtual sample design would be enough to accurately solve the problem. However, the data of real-life samples have significant errors and the construction of the best-approximating plane for such data may depend on the geometrical location of the sample vectors and on the number of these vectors. While the number of virtual samples with error-free spectra is limited by the number of included genomes, the spectra of virtual samples of genomes with errors are linearly independent and their number is unlimited (however, not exceeding the space dimension of 4096, in our case). Below, the above four possible ways of virtual sample design were tested.

Suppose that the number of different genomes in the metagenome is equal to 15 and the average multiplicity of each genome, m, is 10. For each selected value of σ (normal distribution (m, σ)), which produces random deviation of the multiplicity from the average, 100 simulations are performed. In each simulation, a metagenome was constructed of 15 randomly selected genomes and then a certain sample set was generated at the data error of 25% in each selected genome, exactly as it was done in the previous Section. Next, 20 genomes are selected from the library of known genomes. Let 10 of them coincide with 10 genomes of the 15 different genomes of the metagenome, while other 10 genomes are not included in the metagenome. Thus, in each simulation, a total of 25 genomes are present in the samples, 20 of them being added virtually, which gives the accurate number of different genomes equal to 25. If 20 added genomes are subtracted from this number, the final result will be 5 different unknown genomes in the metagenome.

The results of the above calculation for different virtual sample designs are shown in Table 1. It can be seen that the best results were obtained by adding virtual samples with one error-free genome each (case 1). The second-quality result was obtained for virtual samples, each containing one genome with error (case 3). Mixtures of genomes (with errors and without errors) gave worse results, which can be caused by the additional difficulty arising in these cases, namely, by mixing of the genome spectra inside each virtual sample.

**Table 1.** Percentage of correct results for different design of virtual samples. σ is a parameter of normal distribution of the number of genome copies. N is the number of real-life samples taken. The first column shows the types of added virtual samples.

| **Σ** | **4** | **3** |  | **2** |  |
| --- | --- | --- | --- | --- | --- |
| **N** | **20** | **20** | **30** | **30** | **40** |
| **1** | **92** | **89** | **86** | **84** | **80** |
| **2** | **92** | **78** | **81** | **56** | **60** |
| **3** | **90** | **88** | **80** | **82** | **74** |
| **4** | **73** | **61** | **75** | **58** | **66** |

## Discussion

The algorithm proposed in the present work is aimed at calculating or estimating the number of non-identical genomes in the metagenome. We analyze a group of short read sets that are formed as a result of DNA full genome sequencing of the metagenome. Each metagenome sample generates one short read set. The method is basically advantageous in that the algorithm does not require any information about the organisms’ DNAs constituting the metagenome, even though the latter may represent a mixture of different bacteria, protozoa, fungi, or human DNAs.

Although, formally, the proposed approach is applicable to metagenomes containing any number of different genomes, it is obvious that the algorithm effectiveness will decrease with increasing DNA diversity in the metagenome. However, this is true for all methods of metagenome analysis, while for the proposed algorithm, the increase of sequencing accuracy leads to the increase of effectively analyzed metagenome volume. In the case of error-free data, our algorithm provides the true answer, which distinguishes it from numerous algorithms of metagenome analysis, at least from those used for short reads. It should be also noted that the method of using already known genomes (from a certain database), proposed by us, reduces the computational complexity to calculating merely the number of different unknown genomes in the metagenome.

In connection with the above-mentioned drawback, common for all methods of metagenome analysis, various additional techniques for simplifying the data have been proposed in the literature. The method of isolating bacteria using the so-called gnotobiotic animals is of particular interest. According to this method, the microbiota under study is incorporated into sterile mice and diluted several times until only a few microorganism species remain in it [28]. Thus the algorithm proposed by us is also applicable to the most topical studies of human metagenomes.

There also exist natural metagenomes, such as certain metagenomes present in human body, which contain a small number of different genomes. For example, in the urine of some volunteers, a lot of bacteria were found, while in others very few or even only one microorganism was detected [29].

Blood can be a similar object. Previously it was thought to be sterile, but recently microorganisms have been found in it [30]. Perhaps, as in the case of urine, blood may be sterile, but not always. Calculating or at least estimating the number of different microorganisms in blood may become an important part of clinical investigation.

In nature, metagenomes permanently poor in different genomes can be found in extreme environments, e.g., in water sources under extreme conditions. Metagenomes should also be analyzed for sterility control in technical or agricultural systems.

## Methods

The goal of the present study is finding a computer algorithm for calculating the number of different genomes in a metagenome on the basis of its full-genome sequencing, without using additional information about the genomes constituting the metagenome.

The problem to be solved was formulated, the algorithm was proposed, mathematical estimations were obtained for the dependence of the probability of finding the exact number of different genomes on the algorithm parameters, computer simulation to verify the theoretical results was performed.

The computer simulation was performed using 100 bacterial genomes available from site http://www.ncbi.nlm.nih.gov/genomes/lproks.cgi. For each genome, the barcode spectra were calculated based on all possible 6-letter words. Therefore, the BS vector dimension was 4096 and the value of each coordinate was the total number of the corresponding 6-letter word in the genome sequence viewed in both directions ((3’ → 5’ or 5’ → 3’). To simulate the metagenome spectrum, a given number of genomes were randomly (with equal probabilities) sampled and their BS spectra were multiplied by random multiplicities and summed like vectors. Errors simulating sequencing errors were also introduced into the spectra. Then the result of applying the algorithm was compared with the real one. This process was performed repeatedly.

An approach for calculating the number of different genomes based on the principal component method is proposed. All calculations were performed using the MatLab standard functions.

